# Sulforhodamine B and exogenous surfactant effects on alveolar surface tension under acute respiratory distress syndrome conditions

**DOI:** 10.1101/2020.04.08.031526

**Authors:** Tam L. Nguyen, Carrie E. Perlman

**Affiliations:** Department of Biomedical Engineering, Stevens Institute of Technology, Hoboken, NJ

**Keywords:** Acute respiratory distress syndrome, surface tension, surfactant therapy, sulforhodamine B

## Abstract

In the acute respiratory distress syndrome (ARDS), alveolar surface tension, *T*, may be elevated. Elevated *T* should increase ventilation-induced lung injury. Exogenous surfactant therapy, intended to lower *T*, has not reduced mortality. Sulforhodamine B (SRB) might, alternatively, be employed to lower *T*. We test whether substances suspected of elevating *T* in ARDS raise *T* in the lungs and test the abilities of exogenous surfactant and SRB to reduce *T*. In isolated rat lungs, we micropuncture a surface alveolus and instill a solution of a purported *T-*raising substance: control saline, cell debris, secretory phospholipase A_2_ (sPLA_2_), acid or mucins. We test each substance alone; with albumin, to model proteinaceous edema liquid; with albumin and exogenous surfactant; or with albumin and SRB. We determine *T in situ* in the lungs by combining servo-nulling pressure measurement with confocal microscopy, and applying the Laplace relation. With control saline, albumin does not alter *T*, additional surfactant raises *T* and additional SRB lowers *T*. The experimental substances, without or with albumin, raise *T*. Excepting under aspiration conditions, addition of surfactant or SRB lowers *T*. Exogenous surfactant activity is concentration and ventilation dependent. Sulforhodamine B, which could be delivered intravascularly, holds promise as an alternative therapeutic.

**New and Noteworthy:** In the acute respiratory distress syndrome (ARDS), lowering surface tension, *T*, should reduce ventilation injury yet exogenous surfactant has not reduced mortality. We show with direct *T*-determination in isolated lungs that substances suggested to elevate *T* in ARDS indeed raise *T*, and exogenous surfactant reduces *T*. Further, we extend our previous finding that sulforhodamine B (SRB) reduces *T* below normal in healthy lungs and show that SRB, too, reduces *T* under ARDS conditions.

## Introduction

The acute respiratory distress syndrome (ARDS) can occur following a systemic insult, such as sepsis or pancreatitis, or a pulmonary insult, such as pneumonia or aspiration (13, 57). Estimates of ARDS incidence range from 6 to 193 per 100,000 person-years, with the highest estimates from the United States (8, 13, 42, 55), and the novel coronavirus pandemic has no doubt increased incidence. Independent of incidence, reports of in-hospital mortality range from 39 to 56% (8, 42, 49, 55, 60).

In ARDS, alveolar edema liquid impedes gas exchange. Mechanical ventilation supports gas exchange but exacerbates lung injury, which contributes to the high ARDS mortality rate (6, 7, 58). Evidence suggests that lung injury is proportional to alveolar surface tension, *T*, and that *T* may be elevated in ARDS (1, 17, 43, 46, 58).

Various possible causes of elevated *T* in ARDS have been proposed. It has most commonly been suggested that plasma proteins in edema liquid raise *T*. However we and others have shown that plasma proteins do not alter *T* in the presence of an intact surfactant layer, as exists in the lungs (23, 26, 31, 38). Other possible causes include cell debris contamination of the alveolar liquid phase (25); increased activity of phospholipases, particularly secretory phospholipase A_2_ (sPLA_2_) (19, 46); and gastric aspiration, which may lower alveolar pH or wash *T-*raising airway mucins to the alveoli (37).

Given the success of exogenous surfactant therapy in treating neonatal respiratory distress, in which *T* is elevated and residual fetal lung liquid impedes gas exchange (30), it is surprising that surfactant therapy has failed to reduce ARDS mortality (5). The failure may be due to inadequate dosage or delivery strategy (14). However, whether exogenous surfactant can lower *T* in the lungs in the presence of substances suspected of raising *T* in ARDS is not known.

Sulforhodamine B (SRB) could be an alternative *T-*lowering therapeutic for ARDS. Sulforhodamine B is a non-toxic dye that, in combination with albumin, lowers surface tension 27% below normal in healthy lungs (12, 32). Sulforhodamine B could potentially be administered intravascularly, enabling a new delivery route not feasible with exogenous surfactant. However the efficacy of SRB in the presence of substances suspected of raising *T* in ARDS is also not known.

Here, under controlled conditions in isolated rat lungs, we test whether purported *T-*raising substances raise *T* in the lungs and whether exogenous surfactant and SRB can counter the effects of *T*-raising substances. We flood surface alveoli with solutions of potentially *T*-raising substances. We administer each solution alone, with albumin to model the ARDS disease state and with albumin plus exogenous surfactant or SRB. We determine *T in situ* in flooded alveoli.

## Methods

### Isolated lung preparation

We handle all animals in accord with a protocol approved by the Stevens Institute of Technology Institutional Animal Care and Use Committee. As described previously (37), we anesthetize a male or female Sprague-Dawley rat (225-325g, n = 47, Charles River, Wilmington, MA) with 3.5% isoflurane in 100% oxygen. We puncture the heart through the chest wall (21G needle) and euthanize by withdrawal of ∼10 ml blood into a syringe with 1 ml of 1000 units/ml heparin. We cannulate the trachea (blunted 15G needle connected to a stopcock), perform a midline thoracotomy, inflate the lungs with 2 ml air, close the stopcock and excise the heart and lungs. We place the lungs costal surface-upward on the stage of an upright confocal microscope (SP5, Leica Microsystems, Buffalo Grove, IL) and connect an air source and pressure transducer to the tracheal stopcock. Having previously demonstrated that *T* is the same at room temperature as at 37 °C (31), we maintain the lungs at room temperature. We increase transpulmonary pressure, *P*_*L*_, to 30 cmH_2_O and then decrease *P*_*L*_ to a constant 5 cmH_2_O, thus maintaining lung volume above functional residual capacity.

### Base solutions for modeling ARDS

We generate solutions of purported *T-*raising substances, as detailed below, for microinjection into surface alveoli. We use normal saline (NS; pH 5.0) as a control solution and as the solvent for other solutions. For visualization, we include 23 µM fluorescein (00297-17, Cole Parmer, Vernon Hills, IL) in all solutions except Infasurf (35 mg/ml phospholipids, 0.26 mg/ml surfactant protein B and 0.44 mg/ml surfactant protein C; ONY Biotech, Amherst, NY). When we administer Infasurf, we do so as a second injection and include 5 µM sulforhodamine G (SRG, 230561, Sigma Aldrich, St. Louis, MO); we subsequently identify an area in which both fluorescein (first injection) and SRG (second injection) are present. Neither fluorescein nor SRG alters *T* (31, 37).

#### Cell debris

We start by washing the red blood cells (RBCs) of the withdrawn blood. We centrifuge the heparinized blood (5000 x g, 4 °C, 20 min) and discard the supernatant (Fig. 1). We suspend the pellet in pH 7.5 NS to the heparinized-blood volume, centrifuge and discard the supernatant, and repeat the process two more times. Then, to obtain whole-cell debris we homogenize the RBCs or to obtain cell debris fractions we lyse the RBCs.

**Figure 1.**
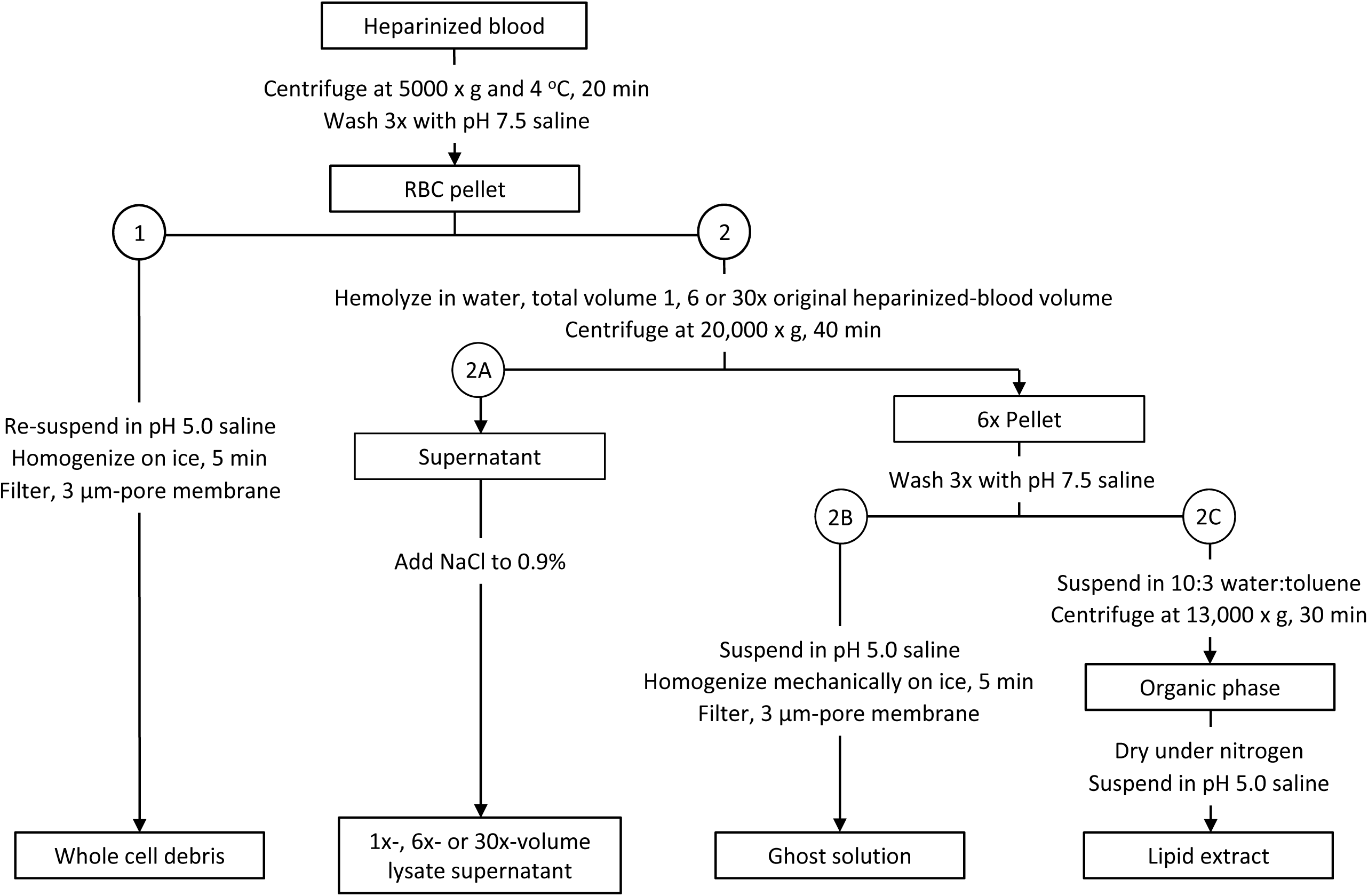
Protocol for obtaining cell debris. Protocol for (1) homogenizing red blood cells (RBCs) to obtaining whole cell debris or (2) lysing RBCs to obtain (2A) lysate supernatant, (2B) ghost solution or (2C) lipid extract.

1.For whole-cell debris, we suspend the pellet in pH 5.0 NS to the heparinized-blood volume. We homogenize the solution on ice (VDI12, VWR, Radnor, PA; maximum speed; 5min) and filter the solution (3-μm pore membrane).

2.For cell debris fractions, we modify the methods of Rosenberg *et al*. (41). We lyse the RBCs by suspending the pellet in 1, 6 or 30x the heparinized-blood volume of Milli-Q-purified water (“water,” pH unadjusted, room temperature, 2 hrs). We centrifuge the lysate (20,000 x g, 4 °C, 40 min), separate the supernatant and pellet and follow one of the three following protocols.

2A. Lysate supernatant. We add NaCl (0.9%) to the supernatant from the 1x-, 6x-, or 30x-volume lysate. The hemoglobin (Hb)-containing solutions are translucent and vary from red (1x) to pink (30x).

2B. Ghost solution. We wash the 6x-lysate pellet by suspending the pellet in the 6x-lysate volume of pH 7.5 NS (more Hb is removed at neutral than acidic pH), centrifuging the suspension (20,000 x g, 4 °C, 40 min) and discarding the Hb-containing supernatant, and repeating the process two more times. We then suspend the pellet in pH 5.0 NS to the 6x-lysate volume, homogenize the solution on ice (maximum speed, 5 min) and filter the solution (3-μm pore membrane). The solution is light pink and cloudy.

2C. Lipid extract of RBC ghost. We wash the 6x-lysate pellet as in 2B and then suspend it in 10:3 water:toluene, with the water volume that of the 6x lysate. To enable phase separation, we vortex the solution briefly, let the solution stand on ice (2 hrs) and then centrifuge the solution (13,000 x g, 4 °C, 30 min). We collect the toluene layer, evaporate the toluene under nitrogen and obtain dry ghost lipids. We weigh some samples, prepared specifically for lipid quantification, on a microbalance. We prepare other samples for alveolar injection by suspending the dry lipids in 100 μl of pH 5.0 NS. The ghost lipid solution is clear, without color.

We quantify Hb concentration, [Hb], of whole-cell debris, lysate supernatants and ghost solution by Drabkin’s assay. We divide each sample for duplicate assay and use bovine methemoglobin (metHb) (H2500, Sigma Aldrich) as a standard. For each solution, we calculate effective 1x [Hb] by multiplying measured [Hb] by dilution factor. For each lysate supernatant, we also prepare a solution of the equivalent concentration of metHb, [metHb].

#### sPLA_2_

We dissolve bovine pancreatic sPLA_2_ IB (P8913, Sigma Aldrich; 0.1 mg/ml) or recombinant rat sPLA_2_ IIA (Uniprot no. P14423 from ELISA kit LS-F23950, LS Bio, Seattle, WA; 2.5 ng/ml) in pH 5.0 NS. These two sPLA_2_ forms are the forms most likely to be present in ARDS (20, 24, 46).

#### Acid and mucins

To test low pH or mucins, we add 0.01 N hydrochloric acid (HCl) or 25 µg/ml porcine gastric mucin (M2378, Sigma Aldrich), respectively, to pH 5.0 NS. Alveolar instillation of this HCl concentration, which has a pH of 1.9, equivalent to that in the stomach (44), represents a worst-case scenario in which un-buffered gastric liquid reaches the alveoli.

### Protocol

We hold *P*_*L*_ at 15 cmH_2_O, puncture a surface alveolus with a glass micropipette filled with a given solution and inject ∼7-10 μl, which floods a group of alveoli. In flooded alveoli, the air-liquid interface forms a meniscus (3, 31).

We determine *T* at the meniscus. We first establish a regular volume history for the lungs. Noting that following alveolar injection of 5% albumin solution *T* is the same after 2 or 60 ventilation cycles between *P*_*L*_ of 5 and 15 cmH_2_O (31), we cycle *P*_*L*_ twice between 5 and 15 cmH_2_O and then hold *P*_*L*_ at 15 cmH_2_O. We determine *T* at this high *P*_*L*_ because we can detect a smaller % change in *T* at higher *P*_*L*_ and because elevated *T* at high *P*_*L*_ is pathophysiologically relevant in that it exacerbates over-expansion induced lung injury (31, 32, 38, 58). We determine alveolar air pressure with a tracheal transducer; liquid pressure in a flooded surface alveolus by servo-nulling measurement (Vista Electronics, Ramona, CA); local three-dimensional meniscus radius in the same alveolus by confocal microscopy; and calculate *T* from the Laplace relation (31, 37).

### Experimental groups

For control pH 5.0 NS and solutions of each potential *T*-raising substance, we test the following groups. (i) Base solution. (ii) To model plasma-protein rich ARDS edema liquid, base solution + 5% bovine serum albumin (A8327, Sigma Aldrich). (iii) To test exogenous surfactant at its full clinical dose, initial administration of base solution + 5% albumin and, ∼1 min later, after cycling twice between *P*_*L*_ of 5 and 15 cmH_2_O to clear unstably-flooded alveoli, subsequent administration to the same region of 100% Infasurf. (iv) When Infasurf does not, after the standard two-cycle ventilation history, lower *T*, we add an additional experiment group in which we assess the effect of further ventilation on exogenous surfactant activity. As in group (iii), we inject base solution + 5% albumin, ventilate twice and inject Infasurf. But then, we cycle 50, rather than 2, additional times between *P*_*L*_ of 5 and 15 cmH_2_O before determining *T*. (v) To test SRB (341738, Sigma Aldrich), single administration of base solution + 5% albumin + 10 nM SRB. Table 1 shows the effects of albumin and SRB on the pH of select solutions.

**Table 1:**
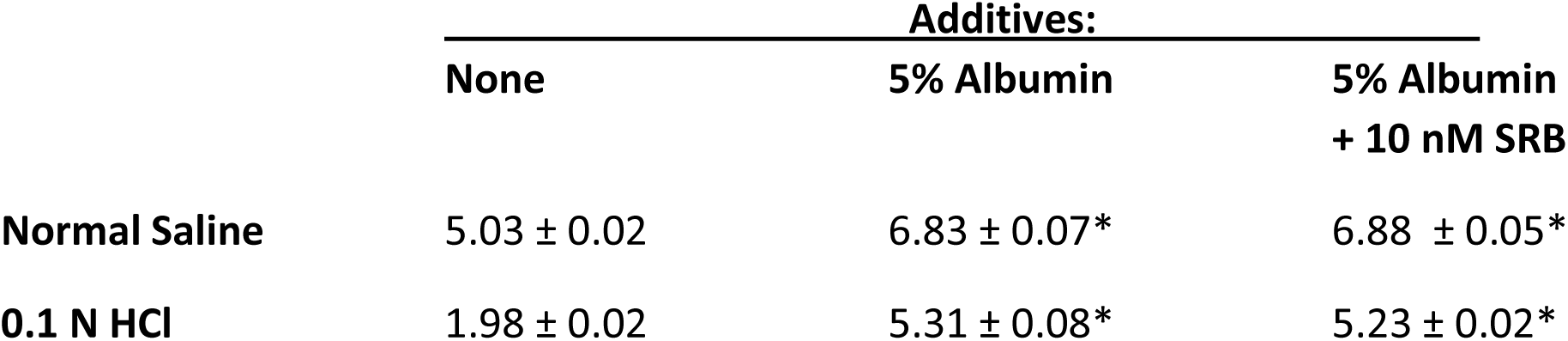
Albumin effects on pH of select solutions. Solution pH measured on bench top at 22°C before injection into alveoli. *p < 0.05 vs. no additives group for same base solution.

In the two-cycle experiments of groups (i-iii) and (v), we flood a region, ventilate twice and determine *T* in a single alveolus in the region, and then repeat the protocol in 3-5 more regions in the same lungs over a period of up to 5 hrs after lung isolation. We test each solution in at least two different rat lungs. Determined *T* value does not correlate with time since lung isolation or rat sex. Each *T* determination requires 10-15 min. As the lungs are held at constant inflation during this time, our measurement is one of static surface tension. However, *T* in the lungs, in contrast to *T* in an *in vitro* surfactometer, remains metastable at the its value at the time at which ventilation is stopped (28, 48). For this reason, the difference observed *in vitro* between static and dynamic *T* may not exist in the lungs. In support of this possibility, *T* determined by our static method in the lungs correlates with degree of dynamic ventilation injury (58).

In the 50-cycle experiments of group (iv), we inject base solution + 5% albumin into a first region, ventilate twice and inject Infasurf into the same region, and then repeat the two-step flooding protocol in a second region. We then administer the 50 ventilation cycles, determine *T* in the second region and determine *T* in the first region. The time from end of ventilation to *T* determination is 15 min longer for the first than the second region. Next, we repeat the entire protocol, with a different solution, for two additional regions. Thus the first set of 50 ventilation cycles applied to the lungs before injection into regions #3 and 4 is an additional protocol step for regions #3 and 4 that is not present for regions #1 and 2. However, we test each solution in two rat lungs – in regions #1 and 2 in one lung and regions #3 and 4 in a second lung. Determined *T* value does not vary with region number.

### Statistical analysis

We analyze data sets (in Figs. 3 and 4 – groupings of 2-5 vertical bars plus normal *T* of the aerated alveolar liquid lining layer, represented by horizontal bar that displays data from Kharge *et al*. (31)) by multiple linear regression. Experimental replicate numbers indicate *T* determinations in individual alveoli. We report group mean ± standard deviation and accept p < 0.05 as significant.

**Figure 2.**
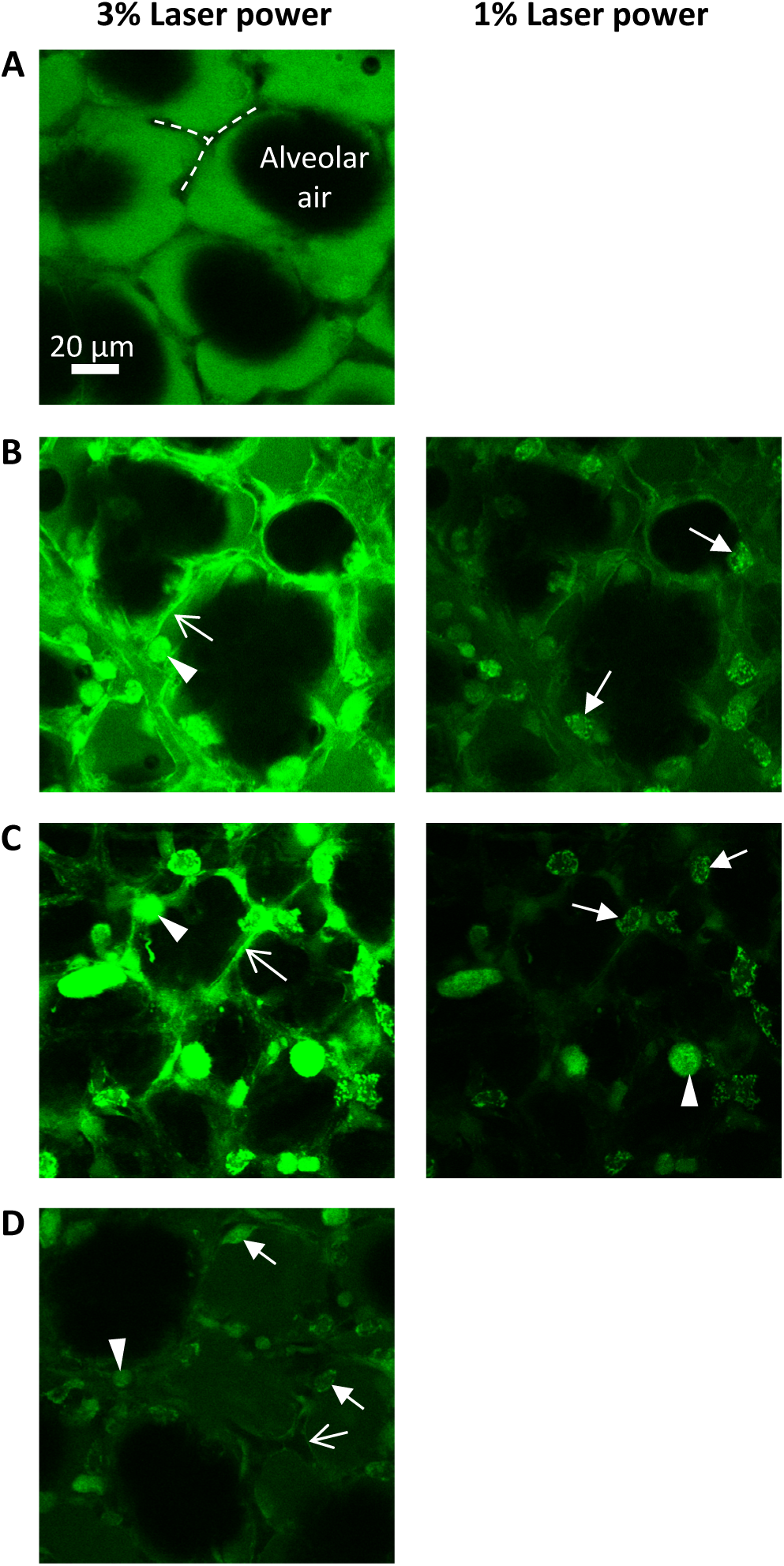
Solution effects on cells. Confocal images of alveoli after flooding with fluorescein (23 µM; green in images) in **(A)** normal saline, **(B)** 0.1 mg/ml secretory phospholipase A_2_ (sPLA_2_) IB, **(C)** 0.01 N hydrochloric acid (HCl) or **(D)** 0.01 N HCl + 5% albumin. In (A), airspace (labeled) and septa (dashed lines) are black. The latter indicates fluorescein exclusion by intact epithelium. In (B-D), fluorescein appears to be concentrated in cells. Morphology suggests that the cell types may include macrophages (arrowheads, round), alveolar epithelial type I cells (open-headed arrows, thin and lining septum) and, likely, alveolar epithelial type II cells (closed-headed arrows, cuboidal), indicating damage to cell membranes and possible macrophage activation. Images (488 nm excitation, indicated laser power, 750 gain) taken ∼2 min after solution injection, at 15 μm sub-pleural depth. Low pH in alveolar liquid quenches fluorescein. Due to saturation in (B) and (C) on left, lower-laser-power replicate images shown on right.

**Figure 3.**
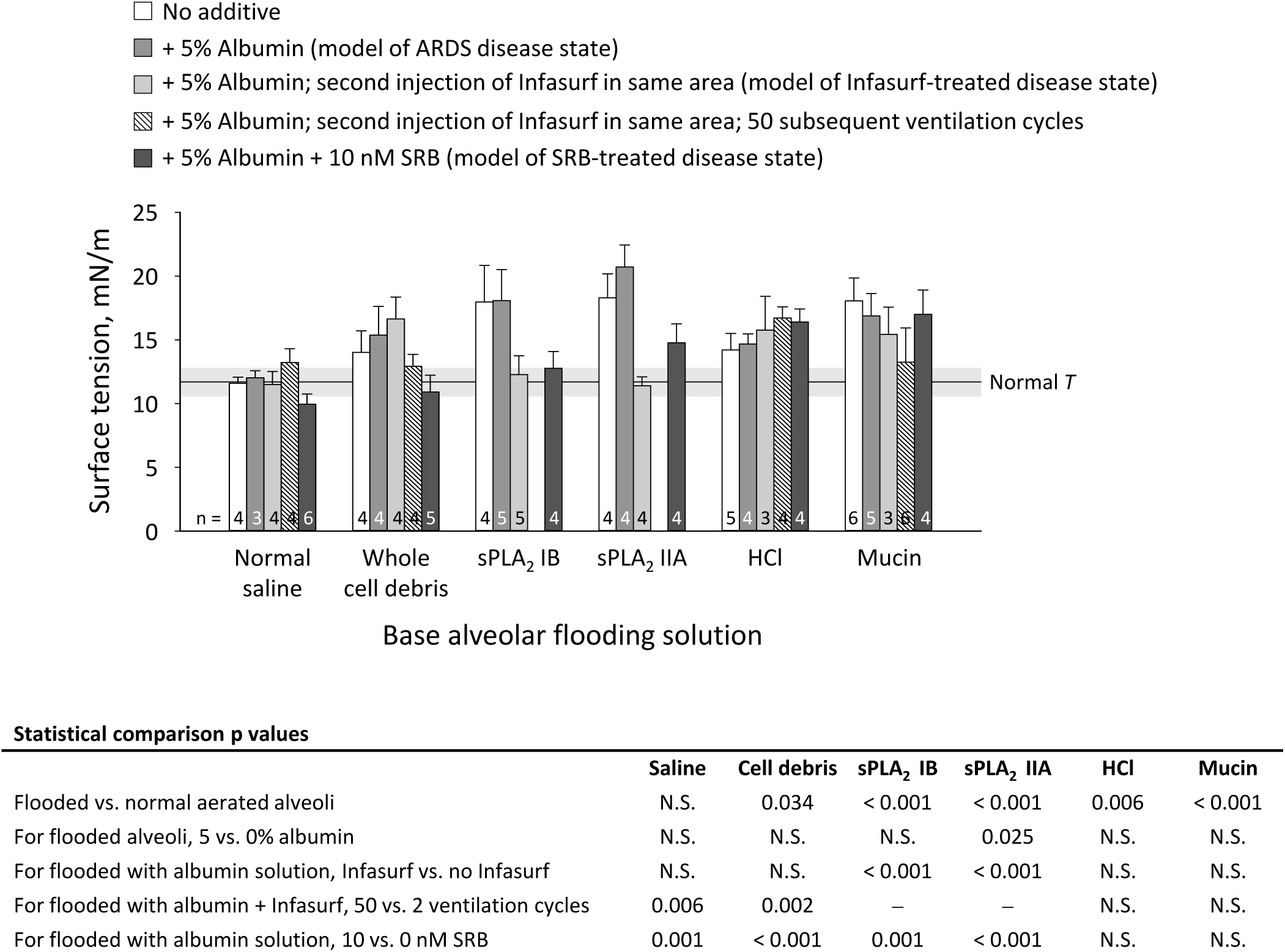
Solution effects on alveolar surface tension, *T*. Base solutions are: normal saline (control solution and solvent of experimental solutions); whole-cell debris, with hemoglobin concentration of heparinized blood; sPLA_2_ IB, 0.1 mg/ml; sPLA_2_ IIA, 2.5 ng/ml; HCl, 0.01 N; and porcine gastric mucin, 25 µg/ml. Additives are as shown. After ventilating twice between transpulmonary pressure, *P*_*L*_, of 5 and 15 cmH_2_O, lungs held at constant *P*_*L*_ of 15 cmH_2_O during 10-15 min period required to determine *T*. For the Infasurf + ventilation group, we apply 50, rather than two, ventilation cycles between *P*_*L*_ of 5 and 15 cmH_2_O. Further details provided in text. Horizontal gray bar shows mean ± standard deviation for *T* of normal liquid lining layer in aerated lungs, also at *P*_*L*_ of 15 cmH_2_O, from (31). Data for “no additive” groups with saline, HCl and mucins are from previous study (37). Statistics: shown on figure; N.S. indicates not significant.

**Figure 4.**
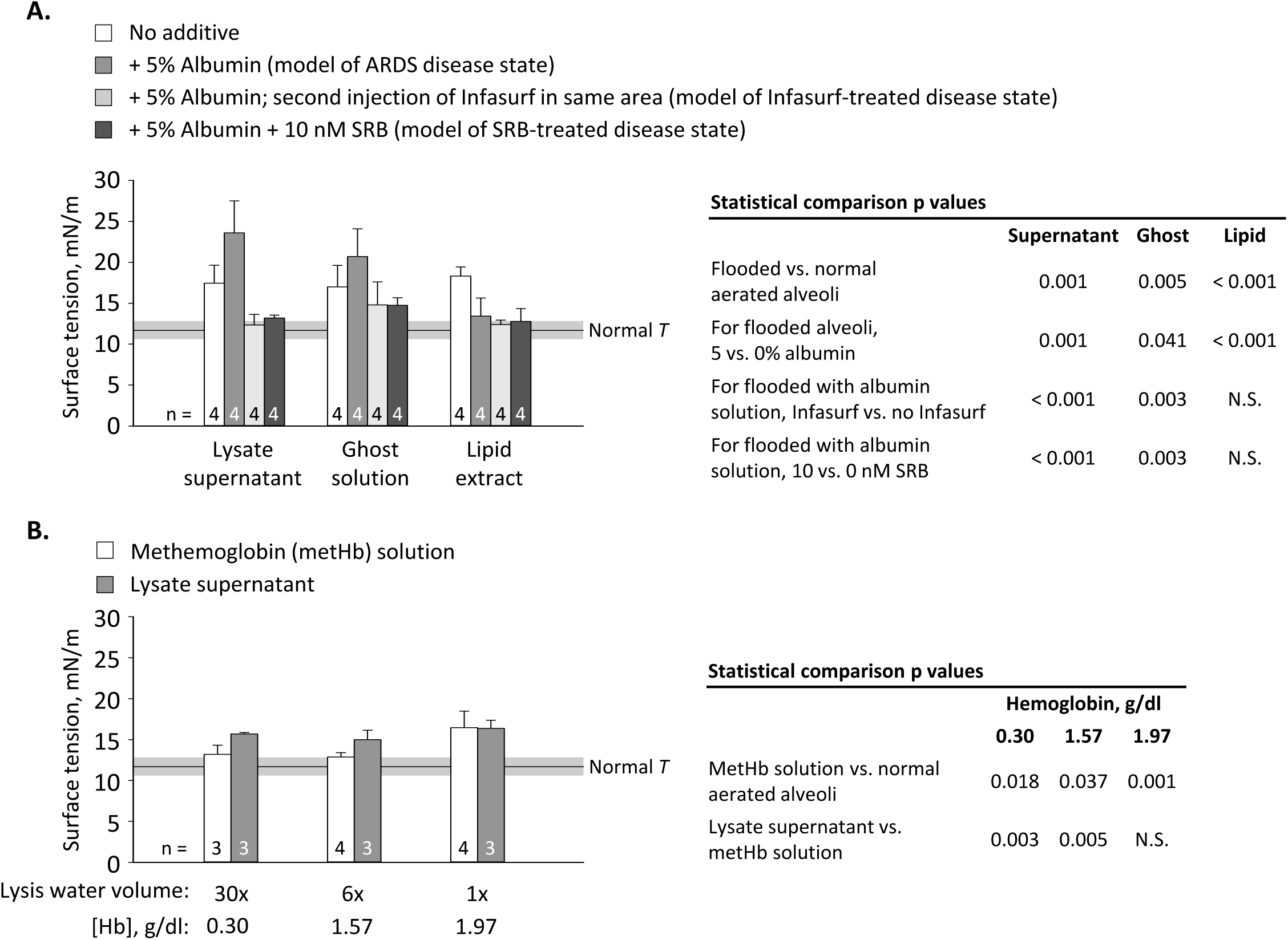
Red blood cell fraction effects on *T*. **(A)** Effects of 6x-lysate fractions, with additions as shown, on *T*. **(B)** Effects of all concentrations of lysate supernatant and solutions with matching concentrations of methemoglobin on *T*. Lysis water volume is multiple of original heparinized-blood volume. Surface tension determination, horizontal bar and statistics as in Fig. 3.

## Results

### Control saline

In all images of injected saline ± albumin, alveolar septa appear black, indicating that fluorescein remains in the alveolar liquid (Fig. 2A). As in previous studies (31, 32, 37), alveolar injection of NS does not alter *T* from normal and addition of albumin has no effect (Fig. 3). Addition of Infasurf (two ventilation cycles) does not alter *T* but Infasurf + 50 ventilation cycles raises *T* above normal. Addition of SRB lowers *T* below normal.

### Cell debris

In all images of injected cell debris ± albumin, septa appear black; fluorescein remains in the alveolar liquid. Homogenized whole-cell debris raises *T* and neither albumin nor Infasurf (two ventilation cycles) has an effect (Fig. 3). However both Infasurf + 50 ventilation cycles and SRB lower *T*.

From the 6x-volume lysate, supernatant, ghost solution and ghost lipid extract fractions raise *T* above normal (Fig. 4A). Inclusion of albumin further raises *T* of lysate supernatant and ghost solution but lowers *T* of lipid extract. Addition of Infasurf (two ventilation cycles) or SRB lowers *T* of lysate supernatant and ghost solution but has no further effect on *T* of lipid extract.

Methemoglobin solutions with concentrations equivalent to the Hb concentrations of all three lysate supernatants (Table 2) raise *T* above normal (Fig. 4B). The 30x- and 6x-lysate supernatants raise *T* more than their equivalent metHb solutions but the 1x-lysate supernatant does not.

**Table 2:** Concentrations of hemoglobin (Hb) and lipids in whole red blood cell (RBC) debris and RBC lysate fractions. Solutions obtained from heparinized blood as depicted in Fig. 1 and described in Methods. Effective 1x concentration is product of measured concentration and dilution factor.

|  | Hb, g/dl<br>(n=3/group) | Lipids, g/dl<br>(n=3/group) | Effective 1x Hb<br>concentration, g/dl |
| --- | --- | --- | --- |
| Whole-cell debris from homogenization | 13.3 ± 0.34 | --- | 13.3 |
| Supernatant from 1x-volume lysate | 1.97 ± 0.27 | --- | 1.97 |
| Supernatant from 6x-volume lysate | 1.57 ± 0.13 | --- | 9.42 |
| Supernatant from 30x-volume lysate | 0.30 ± 0.01 | --- | 9.00 |
| Ghost solution from 6x-volume lysate | 0.12 ± 0.02 | --- | 0.72 |
| Lipid extract from 6x-lysate ghost solution | --- | 0.13 ± 0.02 | --- |

### Phospholipase A_2_

In all images of injected sPLA_2_ IB ± albumin, 1/4 images of sPLA_2_ IIA and 2/4 images of the sPLA_2_ IIA + albumin, fluorescein is concentrated in cells (Fig. 2B). This concentration suggests hydrolysis of cell membranes and possible macrophage activation. Group IB or IIA sPLA_2_ solution raises *T* (Fig. 3). Addition of albumin has no effect on *T* of sPLA_2_ IB solution but further raises *T* of sPLA_2_ IIA solution. For these sPLA_2_ groups without and with albumin, there is no correlation between visual evidence of cell damage and degree of *T* elevation. Addition of Infasurf (two ventilation cycles) or SRB reduces *T*.

### Hydrochloric acid and mucins

Hydrochloric acid solution is the only tested solution that causes regional damage evident in bright-field images as visible discoloration (not shown). Following injection of HCl, fluorescein is concentrated in cells (Fig. 2C, representative of all HCl images) and *T* is elevated (Fig. 3). Addition of albumin buffers HCl and increases pH to 5.3 (Table 1). Nonetheless, following injection of HCl + albumin, fluorescein is concentrated in cells (Fig. 2D, representative of all HCl + albumin images) and *T* remains elevated. Neither Infasurf – even with 50 ventilation cycles – nor SRB lowers *T*.

In all images of injected mucin ± albumin, septa appear black; fluorescein remains in the alveolar liquid. Mucin solution raises *T* and inclusion of albumin has no further effect (Fig. 3). Neither Infasurf – even with 50 ventilation cycles – nor SRB lowers *T*.

## Discussion

We consider, below, the mechanisms through which the substances we test alter *T*.

### Control Saline

Our present results (Fig. 3) agree with our past report that neither albumin nor blood plasma raises *T* (31). Plasma proteins can increase *T* by adsorbing faster than surfactant to a clean interface (with constant area or cyclic 50% area compression) or re-adsorbing faster following layer collapse (caused by 80% area compression *in vitro* or 1-30 cmH_2_O *P*_*L*_ variation in isolated lungs) (23, 26, 31, 56). However, plasma protein addition to the subphase does not alter *T* in the presence of an intact interfacial surfactant layer – *in vitro*, even with 50% area compression, or in isolated lungs, with physiologic *P*_*L*_ (23, 26, 31). With functional native surfactant + albumin, exogenous surfactant + ventilation intriguingly raises *T* but SRB, via a not-yet-determined mechanism, improves the efficacy of native surfactant and lowers *T* below normal.

### Cell debris

In ARDS, including in that caused by novel coronavirus infection, the epithelium is damaged and alveolar hemorrhage is sometimes present (2, 15, 36, 52). For whole-cell debris, Infasurf + 50 ventilation cycles lowers *T* (Fig 3) – interestingly, to a level that appears to be the same as that to which, with NS + albumin (no cell debris), Infasurf + ventilation raises *T*. Sulforhodamine B reduces whole-cell-debris elevated *T* faster than Infasurf, after two ventilation cycles, suggesting action via a different mechanism than exogenous surfactant.

We further determine *T* in 6x-volume lysate fractions in which cell debris is more dilute than in homogenized whole-cell debris (Fig. 4A). The effect of albumin differs between fractions.

Extending the previous finding of Holm and Notter that RBC membrane lipids (extracted from whole sonicated cells) interfere with surfactant adsorption (25), we show lipid extract to interfere with an intact surfactant layer. Red blood cell membrane lipids comprise 54% phospholipid (PL), of which 6.5% is lysophosphatidylcholine (lysoPC) and 4% is disaturated PC (25). Thus membrane lipids are more fluid than surfactant phospholipids and may interfere with the tight packing of surfactant phospholipids. LysoPC, in particular, has been shown to interfere with – suggested to intercalate into – intact surfactant layers (24, 26), though whether our lipid extract contains sufficient lysoPLs to raise *T* (21) is not known. Addition of albumin abolishes the *T*-raising effect of lipid extract. Although we use physiologically-relevant fatty acid-replete albumin, albumin likely sequesters additional lipids.

Ghost solution is a homogenate of a complex mixture of RBC shells comprising cytoskeleton plus plasma membrane; organelles with their own membranes; and some whole RBCs. The presence of whole RBCs is attributable to incomplete hemolysis. During hemolysis in water, quickly-rising osmolarity retards further lysis. Thus the effective 1x [Hb] of each lysate supernatant is less than the [Hb] of whole-cell debris and some Hb remains in the ghost solution (Table 2). However *T* elevation of ghost solution (Fig. 4A) is greater than any *T-*elevation its low concentration of Hb could be expected to cause (Fig. 4B), suggesting that Hb is not the main *T-*raising component of ghost solution. Rather, membranes might be the *T-*raising component of ghost solution. This possibility is supported by the lack of effect of albumin – membranes are too large to be sequestered by albumin.

Hemoglobin on its own raises *T* in a dose-dependent fashion (Fig. 4B). One possible mechanism through which Hb might raise *T* is ionic interaction with the negatively-charged heads of surfactant phospholipids, which could destabilize the surfactant layer (50). Previously, Hb concentrations of 2.5-20 g/dl were tested during co-adsorption with surfactant and shown to raise *T* (25). Here, we extend the finding to lower Hb concentrations and an intact surfactant layer.

The 30x- and 6x-lysate supernatants raise *T* more than their corresponding metHb solutions (Fig. 4B). The difference could be due to the presence of Fe^2+^-carrying fresh Hb in lysate supernatant vs. Fe^3+^-carrying metHb in hemoglobin-only solution or due to additional *T-*raising substance(s), beyond Hb, present in lysate supernatant. That fresh Hb is less likely than metHb to disrupt the surfactant layer (50), suggests that the difference is most likely explained by additional *T-*raising substances in lysate supernatant. The identity of those *T-*raising substances and the mechanism through which albumin further raises *T* (Fig. 4A) remain to be determined. The 1x-lysate supernatant has a higher [Hb] and should have a higher concentration of other *T-*raising substances than the 30x- and 6x-lysate supernatants but does not raise *T* more than its corresponding metHb solution. The higher [Hb] has a strong effect on *T* and may mask the effect of other substances.

Sulforhodamine B, which rapidly normalizes *T* of whole-cell debris (Fig. 3), also rapidly normalizes *T* of cell lysate fractions (Fig. 4A). The effect of Infasurf is more complex. With less-concentrated cell-lysate fractions (Fig. 4A), Infasurf immediately normalizes *T*. With more-concentrated whole-cell debris (Fig. 3), ventilation is required for Infasurf to normalize *T*. These results suggest that cell debris-surfactant interaction is concentration dependent and support the prior finding (e.g., 35, 53) that ventilation promotes surfactant activity.

### Secretory PLA_2_

How elevated sPLA_2_ activity may elevate *T* in ARDS has not been fully explained. First, the mechanism could be surfactant phospholipid depletion, but >80% hydrolysis might be required (21, 24). As the hydrolysis product lysoPL associates with, but does not completely compose, small aggregates and small aggregates contain 72% of ARDS bronchoalveolar lavage fluid (BALF) phospholipids (45, 46), this threshold is not likely met. Second, hydrolysis of plasma membranes (Fig. 2B) might release *T-*elevating cell debris. Third, the surfactant- and plasma membrane-hydrolysis products lysoPL and fatty acid (FA) could disrupt surfactant phospholipid packing. LysoPLs, at 10-20% hydrolysis, raise *T* of an intact surfactant layer, raise *T* following co-adsorption with surfactant and, subsequent to co-adsorption, raise minimum *T, T*_*MIN*_, following cyclic 50% area compression (21, 24, 26). While lysoPLs often comprise <10% of ARDS BALF phospholipids, lysoPC in one case comprised ∼18% (45, 46). Fatty acids, at 10-20% hydrolysis, raise *T* during co-adsorption with surfactant and subsequently, albeit inconsistently and to a lesser degree than lysoPLs, raise *T*_*MIN*_ following cyclic 50% area compression (21, 24). Fourth, the activity of sPLA_2_ IIA, which cleaves phosphatidylglycerol (PG) and phosphatidylethanolamine (PE) but not PC, is elevated in ARDS BALF and the fraction of PG is reduced (22, 45, 46). Degradation of PG, which interacts with surfactant protein B, may have an outsized effect on *T* (46).

We find that albumin does not alter or increases the sPLA_2_ effect on *T* (Fig. 3). These results fit within a wide range of reports of condition-specific and interdependent sPLA_2_-product and albumin effects on sPLA_2_ activity. Plucthun and Dennis (39) showed that FAs stimulate sPLA_2_ IB hydrolysis of PC/ or PE/Triton-X 100 micelles. Also for sPLA_2_ IB, Conricode and Ochs (9) showed that cleavage products inhibit and ≤1% albumin stimulates hydrolysis of PC liposomes; and <0.1% albumin stimulates and 0.1-0.3% albumin inhibits hydrolysis of PC/cholate micelles. (Conricode and Ochs did not investigate ∼5% albumin.) Conricode and Ochs surmised that, as FAs stimulate sPLA_2_ IB, likely by remaining at and lending a negative charge to the lipid-water interface, (i) it must be lysoPLs, more of which dissociate, that inhibit PC liposome hydrolysis and (ii) albumin must sequester lysoPLs. Conricode and Ochs further speculated that different lipid-water interface charges, which underlie membrane affinities for FAs, explain the different albumin effects under different conditions. Our observation that albumin does not alter the sPLA_2_ IB effect on *T* suggests that, for native surfactant, product inhibition of sPLA_2_ IB may be minimal. Although we are not aware of product stimulation/inhibition data for sPLA_2_ IIA, our observation that albumin tends to increase *T* suggests product inhibition and albumin sequestration of product.

As noted above, sPLA_2_ likely cleaves surfactant and plasma-membrane PLs. Infasurf immediately and fully reverses *T* elevation by either sPLA_2_ IB or IIA. That supplementation of degraded native surfactant normalizes *T* again suggests that surfactant activity is concentration dependent. Sulforhodamine B fully reverses *T* elevation by sPLA_2_ IB and partially but significantly reverses *T* elevation by sPLA_2_ IIA. As SRB acts in conjunction with albumin and native surfactant, the ability of SRB to lower *T* indicates that significant functional native surfactant remains present.

### Acid aspiration

In aspiration, pH 1.6-2.3 gastric contents (44) enter the airways and mix with airway liquid that should buffer pH but contribute *T-*raising mucins (33, 37). If there is little mixture with airway liquid then low pH liquid, which should aggregate mucins and block their *T-*raising effect (27, 37), may reach the alveolus. Acid might raise *T* by direct effect on surfactant or indirect effect on the alveolar wall, the latter scenario leading to cell debris contamination of the edema liquid. That Infasurf counters *T* elevation by RBC debris but not acid suggests that cell debris is not the cause of acid-induced *T* elevation; acid may, rather, directly alter lung surfactant. This conclusion contradicts a prior finding of pH of 2 not altering surfactant function *in vitro* (18) but, as mentioned above, surfactant behaves differently in the lungs than *in vitro* (28, 48). If there is significant mixture of aspirate with airway liquid, then it may be a higher-pH liquid that reaches the alveolus. If pH exceeds 4, mucins should be un-aggregated (27) and, upon reaching the alveoli, could potentially raise *T*. The possibility of a detrimental role for mucins stems from our prior finding that tracheal saline instillation washes mucins to the alveoli and raises alveolar *T* 42% above normal (37). However, tracheal saline instillation does not cause lung injury and we determined *T* in that study after two ventilation cycles. Additional ventilation might enable native surfactant to counter the effect of mucins. The pathophysiologic significance of mucin-elevated *T* is not known.

That albumin-buffered HCl, with a pH of 5.3 (Table 1), injures the epithelium (Fig. 2D) is surprising. We speculate that solution pH may decrease upon injection into the alveolus, where phospholipid head group phosphates may induce H^+^ dissociation from albumin.

Neither Infasurf nor SRB reduces *T* elevation by acid + albumin. The inefficacy of SRB against HCl + albumin could be attributable to a pH effect on SRB-albumin interaction. Albumin Sudlow site I attracts the xanthene rings of SRB (34). As discussed above, alveolar injection may reduce the pH of HCl + albumin solution. And pH <4.3 causes compaction of albumin (4, 11), which might interfere with albumin-SRB binding. Alternatively, SRB likely responds to pH similarly to rhodamine WT, which below pH 4.7 loses a negative charge and becomes a zwitterion (54). The lost charge, which is not on the xanthene rings, might alter interaction of an albumin-SRB complex with surfactant.

Neither does Infasurf or SRB reduce *T* elevation by mucins, though Infasurf + ventilation appears close to having an effect. We speculate that the large, non-glycosylated hydrophobic domains of mucins (47, 51) interfere with surfactant by hydrophobic interaction. For SRB there might, similarly, be a hydrophobic interaction between mucins and its xanthene rings (34).

### Study limitations

Our study includes multiple limitations. First, we do not test all purported mechanisms of *T* elevation in ARDS. We do not investigate possible cholesterol contamination of the surfactant layer (16, 35). However we previously showed that cholesterol-containing blood plasma does not alter *T* in the lungs (31). Another possible mechanism is surfactant damage by leukocyte- or macrophage-released reactive oxygen species (ROS). As discussed elsewhere (38), the principal effect of phospholipid oxidation is likely due to generation of lysophospholipids (40). As Infasurf and SRB reverse sPLA_2_-elevated *T*, which is likely attributable to lysophospholipid generation, exogenous surfactant and SRB would both likely reduce *T* elevated by phospholipid oxidation. Oxidation of surfactant protein B (SP-B), however, may also contribute to *T* elevation in ARDS (40); whether oxidation of SP-B raises *T* of an already-established intact surfactant layer and whether exogenous surfactant or SRB would reduce *T* in such a scenario remains to be determined. Second, we do not assess alveolar diminishment or collapse that might rupture the surfactant layer and allow edema liquid components to raise *T* by adsorption. Further, surfactant secretion and recycling are continuous processes in the alveolus, including in isolated lungs (29). We have not determined whether *T-*raising substances act by interfering with secretion or recycling. Finally, for the substances we test, we do not know the pathophysiologic concentration in ARDS edema liquid. We select concentrations that raise *T* in the lungs to levels comparable to those in other models of lung injury (31, 37, 59). To determine whether SRB can reduce *T* under ARDS conditions, testing in animal models of ARDS will be required.

### Conclusion

We show that cell debris, sPLA_2_, acid and mucins raise *T* of an intact surfactant layer. In the ∼12% of ARDS cases caused by gastric aspiration (13), neither exogenous surfactant nor SRB appears likely to reduce *T*. In ARDS due to other causes, both surfactant therapy and SRB show potential for lowering alveolar *T*. For exogenous surfactant, this study extends prior *in vitro* findings to show ventilation- and concentration-dependent efficacy in the presence of an initially-intact surfactant layer. For SRB, this study extends our previous findings from healthy lungs to show efficacy under ARDS conditions.

Either surfactant therapy or SRB could be administered by tracheal instillation or nebulization. However delivery via the trachea to injured lung regions is a challenge (10, 14). Given the elevated barrier permeability of injured lung regions in ARDS, SRB might alternatively be delivered intravenously. This consideration raises the possibility of a new treatment modality for ARDS.

## Acknowledgement

We are grateful to Dr. Edmund Egan, of ONY Biotech, for donating the Infasurf for this study.

